# Born to be asocial: newly-hatched tortoises spontaneously avoid unfamiliar individuals

**DOI:** 10.1101/152314

**Authors:** Elisabetta Versace, Silvia Damini, Matteo Caffini, Gionata Stancher

## Abstract

Individual recognition is important for modulating social interactions but it is not clear to what extent this ability depends on experience gained through repeated interactions with different individuals. In wild tortoises, evidence of social interactions is limited to behaviours performed years after hatching, in the context of mating. To investigate the presence of abilities of individual recognition at the onset of life in tortoises, we used hatchlings of two species (*Testudo marginata*, *Testudo graeca*) reared with a single conspecific as unique social experience. When located in a novel environment together with the familiar conspecific, tortoises reached the average distance expected by random trajectories. On the contrary, tortoises tested with an unfamiliar conspecific first explored the mate, then actively kept a distance significantly larger than expected by chance. These results show spontaneous abilities of individual recognition in a non-social species at the onset of life, and active avoidance of unfamiliar conspecifics. We suggest that this predisposed behaviour might be adaptive for young tortoises’ dispersal.

## BACKGROUND

Individual recognition requires to identify a specific organism according to its distinctive features. This capacity is important to social responses in long-term social contexts [reviewed in, 1,2]: individual recognition of mate and kin can induce a closer relationship to familiar individuals, in neighbour-stranger discrimination it modifies responsiveness and aggression towards neighbours compared to strangers, in dominance hierarchies triggers differential responses depending on the relationship with the identified individual. Precocial avian species provide evidence of individual recognition at the onset of life, since newly-hatched birds are adapted to recognize and follow social partners after a brief exposure, through the mechanism of filial imprinting [3–6]. It is not clear whether, beside filial imprinting, individual recognition is available at the onset of life, and whether this ability is present in species with limited social habits.

We addressed these issues investigating newly-hatched tortoises, precocious animals that are known as non-social. In fact, tortoises do not exhibit post-hatching parental care, mate promiscuously and do not form pair bonds or cohesive social groups [7,8]. In wild tortoises, evidence of social interactions is limited to behaviours that are performed when sexual maturity is reached, years after hatching, such as courtship, mounting and nesting [see for instance, 9–12]. In captivity, adult tortoises housed together with conspecifics show a capacity to follow the gaze of conspecifics [13] and to learn from the actions of other individuals [14], suggesting that these animals possess capacities to deal with social partners. It is not clear, though, whether these abilities emerge in captivity as a result of repeated interactions or constitute the spontaneous behavioural repertoire of tortoises. Moreover, it is not clear whether tortoises are capable of individual recognition and if this capacity is present at the onset of life.

To investigate the presence of spontaneous abilities of individual recognition, we used hatchlings of two tortoise species (*Testudo marginata*, *Testudo graeca*). After hatching in individual compartments, we raised young tortoises with a single conspecific as unique social experience, before testing them in a novel environment together with a familiar tortoise (familiar condition) or an unfamiliar tortoise (stranger condition). We expected to observe different responses to familiar *vs.* unfamiliar individuals only if tortoises were capable of individual recognition. We observed a different behaviour in familiar and stranger pairs, showing capabilities of spontaneous individual recognition at the onset of life in these species. By using as experimental setup a circumscribed arena, we were able to compare average pairwise distance with that expected with random trajectories and to evaluate whether tortoises stayed closer of further than expected by chance. While familiar tortoises progressively reached the chance distance, unfamiliar tortoises initially explored the unfamiliar mate, then actively kept a distance larger than expected by chance.

## METHODS

### Subjects and rearing conditions

We observed 26 newly hatched tortoises of two species: 14 *Testudo graeca* and 12 *Testudo marginata* individuals. Tortoises were about one-month old (24-43 days of age, average 27 days) at the moment of the test. Eggs laid on the ground by tortoises were collected by the experimenters and incubated in darkness at 31 °C +/-2 °C. Tortoises hatched in individual compartments and did not see any conspecific for about 10 days (2-20 days, average 10 days) before being paired with a tortoise of the same species. Tortoises were fed with green leaves and hydrated at least twice daily. Each pair was housed in a square-shaped arena (20x20x12 cm) with the bottom covered with soil, leaves and straw. Before the test, subjects did not see any other tortoise and did not interact with the experimenters.

### Experimental apparatus

As experimental apparatus, we used a circular arena (Ø 25 cm, 10 cm high, Fig. 1a) with the bottom covered with wet sand (0.5 cm). A Windows LifeCam© camera located on top of the centre of the arena recorded tortoises’ behaviour.

**Figure 1.**
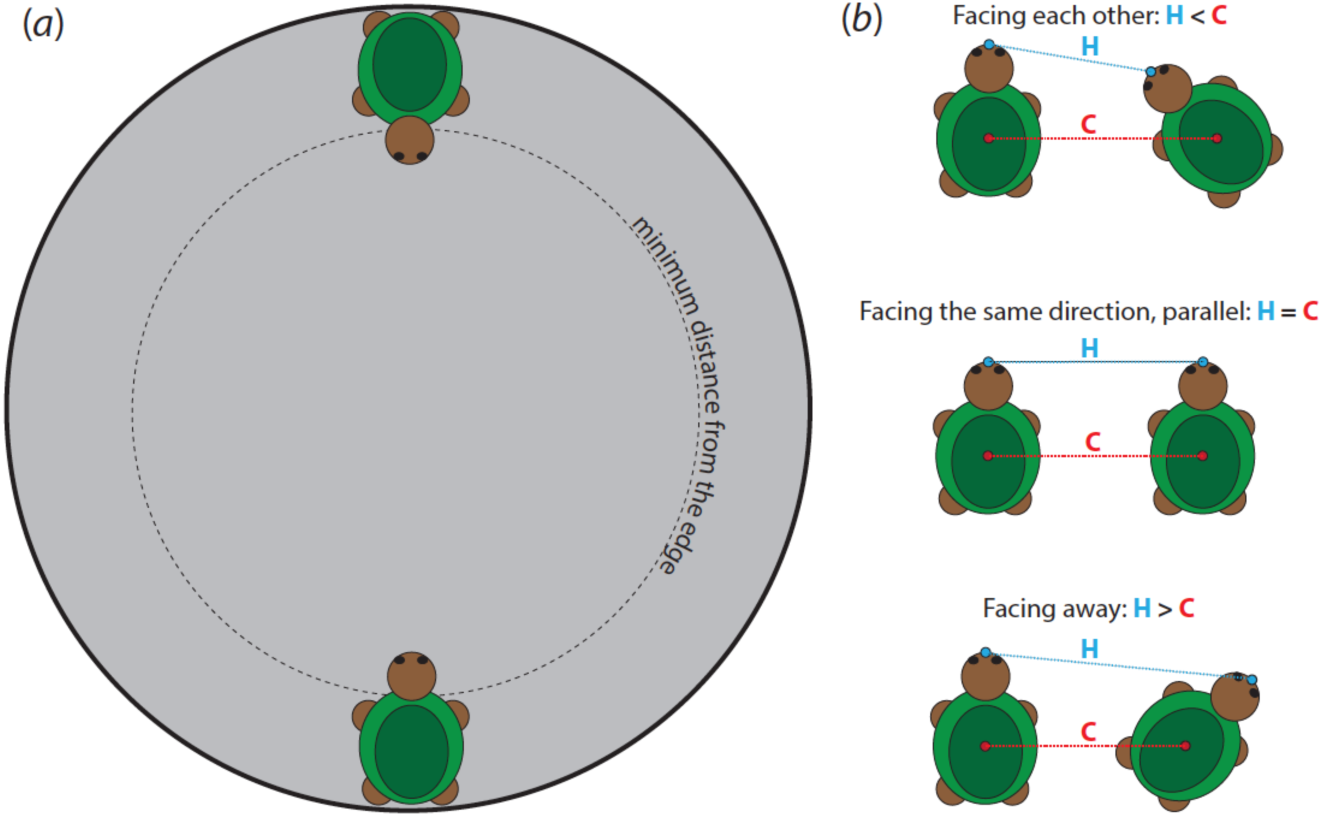
(a) Schematic representation of the experimental apparatus with a pair located in the starting position. (b) Pairwise distance between heads (H) and carapaces (C). When H<C tortoises face each other, when H=C tortoises are parallel, facing the same direction, when H>C tortoises face away from each other.

### Procedure

We first familiarized tortoises with a mate by keeping a pair in the same enclosure for about 22 days (17-33 days, average 22.5 days). Tortoises were then tested in pairs of the same species. Each tortoise was tested once or twice. The list of experimental pairs is shown in Table 1.

**Table 1.** List of experimental pairs by Species and Condition.

| Species | Condition | Pair |
| --- | --- | --- |
| *T. graeca* | Familiar | 15-16 |
| *T. graeca* | Familiar | 17-21 |
| *T. graeca* | Stranger | 11-13 |
| *T. graeca* | Stranger | 12-14 |
| *T. graeca* | Stranger | 23-31 |
| *T. graeca* | Stranger | 25-42 |
| *T. graeca* | Stranger | 37-24 |
| *T. marginata* | Familiar | 22-28 |
| *T. marginata* | Familiar | 3-4 |
| *T. marginata* | Familiar | 5-6 |
| *T. marginata* | Familiar | 7-8 |
| *T. marginata* | Stranger | 1-10 |
| *T. marginata* | Stranger | 9-2 |
| *T. graeca* | Familiar | 11-12 |
| *T. graeca* | Familiar | 13-14 |
| *T. graeca* | Familiar | 23-24 |
| *T. graeca* | Familiar | 25-31 |
| *T. graeca* | Familiar | 37-42 |
| *T. graeca* | Stranger | 16-17 |
| *T. marginata* | Familiar | 1-2 |
| *T. marginata* | Familiar | 9-10 |
| *T. marginata* | Stranger | 3-6 |
| *T. marginata* | Stranger | 5-8 |
| *T. marginata* | Stranger | 7-4 |

Before the beginning of the experimental session, we regulated the external temperature of the subjects under a light, to make sure the temperature measured on the top of the carapace differed less than 2 °C between tested individuals. We measured the carapace temperature with an infrared thermometer. Immediately before the test, each individual was isolated for 5 minutes in an opaque box. Subsequently, the experimental subjects were located in front of each other (a familiar or a stranger tortoise, according to the experimental condition), facing the centre of the arena, at diametrically opposed positions (the furthest possible distance within the arena). The behaviour was recorded for 15 minutes from the moment in which one tortoise of the tested pair moved the first step. We defined as the first step a movement of at least one leg that displaced the carapace. If both tortoises did not move for more than 10 minutes, the session was aborted and repeated the subsequent day. To score the behaviour of tortoises, we extracted one frame every 20 seconds (3 frames per minute) and used ImageJ [15] to identify the location of the centre of the carapace and the tip of the head of each tortoise in all frames.

For each pair we calculated, for 5 consecutive periods of three minutes (15 minutes overall), the distance between the centroids of the carapaces (C), the distance between the tip of the heads (H) in centimetres and the difference between these measures (H-C). The distance between centroids of the carapaces provided and index of proximity irrespective of the relative orientation of the tortoises. The difference between the distance of the centroids and the distance of the heads provided an index of the relative orientation of the subjects: negative values indicate that tortoises are facing each other, positive values that tortoises are facing away from each other (see Figure 1b).

### Identification of the distance expected with random trajectories

To calculate the chance pairwise distance between tortoise centroids (carapace centroid) and between tortoise heads (tip of the head) we implemented a Fermi-like estimation method. Tortoises were simulated as circles with radius 1.82 cm with a circular head of radius 0.5 cm. To obtain the chance carapace distance, we simulated the random positions of 2.5x10^7^ pairs of tortoises uniformly distributed within an arena of radius 12.5 cm (the same size used in the experiments), and calculated their Euclidean pairwise distances. For the chance head distance, we first simulated the position of the carapace centroid, then simulated a random orientation of the head across 360° around the centroid, and finally moved 1 cm away from the edge of the carapace. To avoid overlaps between individuals, we dropped pairs closer than twice the radius of an average tortoise (carapace). To exclude overlaps with the edge of the arena, we dropped positions of individuals closer to the edge than the radius of the tortoise.

We repeated the simulation 20 times and obtained a chance carapace distance of 8.84 cm and a chance head distance that converged to 8.91 cm. All simulations were run with a custom MATLAB code.

### Data analysis

We investigated the relationship between the dependent variables (Overall distance run, Distance between centroids, Distance between heads) and the independent variables Condition (familiar *vs.* stranger), Species (*T. marginata*, *T. graeca*) and Time (1-3, 4-6, 7-9, 10-12, 13-15 minute) as fixed effects and Pair as random effect. Initially, we included the full set of explanatory variables and interactions and progressively found the minimum adequate model using F-tests for models fitted using maximum likelihood. For the variables Distance between centroids and Distance between heads, that showed a significant interaction between Condition and Time, we run post-hoc analyses of variance for the first and second half of the test separately. We used one sample t-tests against the chance level obtained in the simulation to check for significant departures of the distance between centroids and of the distance between heads, independent t-test to compare distance between conditions at specific time points and paired t-test to compare distance within condition at two time points. Significance was set at p<0.05. Analysis were conducted with R (version 3.2.1), for model-fitting we used the nlme package.

### Ethics

All experiments comply with the current Italian and European Community laws for the ethical treatment of animals. The experimental procedures were approved by the Ethical Committee of the Fondazione Museo Civico Rovereto (Italy).

## RESULTS

### Distance between carapace centroids

Distance between carapace centroids of the tortoises was analysed with Condition, Species and Time as fixed effects and Pair as random effect, using a linear mixed effect model fitted using maximum likelihood. The minimum adequate model included Condition, Time and their interaction: Condition (F_1,22_=0.444, p=0.512), Time (F_4,88_=7.694, p<0.0001), Condition x Time (F_4,88_=3.003, p=0.023), see Figure 2A.

**Figure 2.**
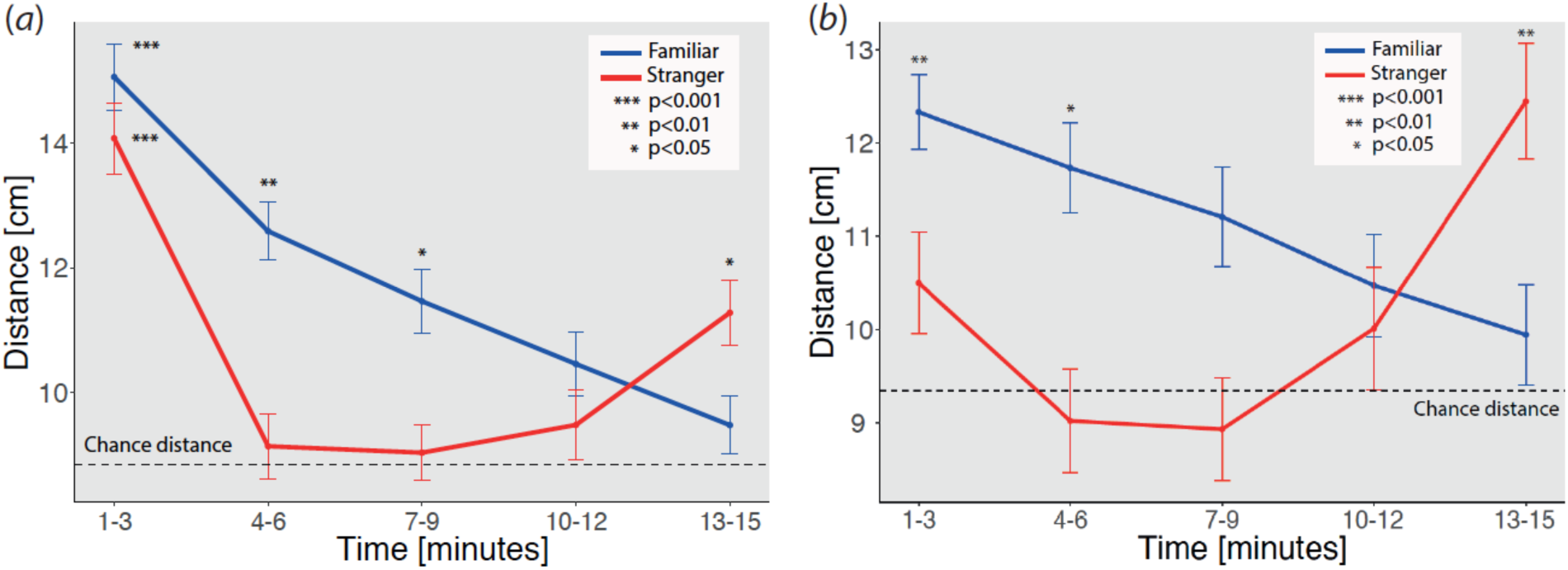
Average distance in centimetres by Condition in Time. (*a*) Average carapace distance. (*b*) Average head distance. The dashed line indicates the chance distance calculated with Fermi-like simulations. Significance against the chance level is indicated by asterisks: *** p<0.001, ** < 0.01, * p<0.05.

As post-hoc tests, we run one analysis of variance for the first and one analysis of variance for the last half of the experiment, using Condition as between factor and Time as within factor. In the first part of the experiment there was no significant main effect of Condition (F_1,22_=0.411, p=0.528) a significant main effect of Time (F_2,44_=7.50, p=0.002) and no significant interaction (F_2,44_=1.559, p=0.222). In the second half of the experiment there was no significant main effect (Condition F_1,22_=2.967, p=0.100; Time F_2,44_=1.730, p=0.189) but a significant interaction Condition x Time (F_2,44_=3.613, p=0.035). As reported in Table 2, while in the familiar condition tortoises progressively approached the chance level distance, in the stranger condition tortoises initially approached the other subject and then moved significantly further than the chance distance. In all time points and conditions, carapace pairs were significantly closer than the average starting distance (20.74 cm).

**Table 2.** Average carapace distance in centimetres, t and p level for one-sample t-tests against the chance carapace distance (8.84 cm) obtained by simulated data.

| Condition | Time [minutes] | Carapace distance [cm] | t | p |
| --- | --- | --- | --- | --- |
| Familiar | 1-3 | 15.07 | 5.068 | <0.001 |
| Familiar | 4-6 | 12.59 | 3.252 | <0.01 |
| Familiar | 7-9 | 11.46 | 2.179 | 0.050 |
| Familiar | 10-12 | 10.46 | 1.335 | 0.207 |
| Familiar | 13-15 | 9.41 | 0.650 | 0.528 |
| Stranger | 1-3 | 14.08 | 5.592 | <0.001 |
| Stranger | 4-6 | 9.13 | 0.244 | 0.812 |
| Stranger | 7-9 | 9.03 | 0.228 | 0.824 |
| Stranger | 10-12 | 9.48 | 0.605 | 0.559 |
| Stranger | 13-15 | 11.28 | 3.006 | 0.013 |

### Distance between heads

Distance between head tips was analysed with Condition, Species and Time as fixed effects and Pair as random effect, using a linear mixed effect model fitted using maximum likelihood. The minimum adequate model included Time as only significant main effect Time (F_4,92_=3.428, p=0.015), see Figure 2B. As post-hoc tests we run one analysis of variance for the first and one analysis of variance for the last half of the experiment, using Condition as between factor and Time as within factor. In the first part of the experiment there was no significant main effect of Condition (F_1,22_=1.730, p=0.202) of Time (F_2,44_=0.712, p=0.49) and no significant interaction (F_2,44_=0.201 p=0.819). In the second half of the experiment there was no significant main effect (Condition F_1,22_=2.485, p<0.001; Time F_2,44_=0.702, p=0.501) but a significant interaction Condition x Time (F_2,44_=4.267, p=0.020). As reported in Table 3, while in the familiar condition tortoises progressively approached the chance level distance, in the stranger condition tortoises initially approached the other subject and then moved significantly further than the chance distance. In all time points and conditions, head pairs were significantly closer than the average starting distance (14.87 cm).

**Table 3.** Average head distance in centimetres, t and p level for one-sample t-tests against the chance carapace distance (9.35 cm) obtained by simulated data, grouped by Condition and Time point (in minutes).

| Condition | Time<br>[minutes] | Head<br>distance<br>[cm] | t | p |
| --- | --- | --- | --- | --- |
| Familiar | 1-3 | 12.33 | 3.890 | 0.002 |
| Familiar | 4-6 | 11.73 | 2.36 | 0.036 |
| Familiar | 7-9 | 11.21 | 1.670 | 0.115 |
| Familiar | 10-12 | 10.47 | 1.047 | 0.316 |
| Familiar | 13-15 | 9.849 | 0.498 | 0.627 |
| Stranger | 1-3 | 10.50 | 1.134 | 0.283 |
| Stranger | 4-6 | 9.023 | -0.268 | 0.794 |
| Stranger | 7-9 | 8.93 | -0.418 | 0.684 |
| Stranger | 10-12 | 10.01 | 0.553 | 0.592 |
| Stranger | 13-15 | 12.44 | 3.261 | 0.009 |

### Facing orientation

The facing orientation, measured as difference between carapace distance and head tips was analysed with Condition, Species and Time as fixed effects and Pair as random effect, using a linear mixed effect model fitted using maximum likelihood. We did not observe significant effects or interactions. However, limiting the analysis to the first two time-points we observed a significant effect of Time (F_1,22_=16.22, p<0.001) no significant effect of Condition (F_1,22_=1.436, p=0.24) and a significant interaction Condition x Time (F_1,22_=5.521, p=0.028), see Figure 3. As showed by one-sample t-tests against the chance level (0), tortoises showed a different propensity to face each other during the test: at time point 1-3 minutes, tortoises of both conditions exhibited a significant preference to face each other (familiar: t_12_=−5.146, p<0.001; stranger: t_10_=−8.858, p<0.001), while at time point 12-15 minutes tortoises in the stranger condition had a significant preference to face in different directions (t_12_=3.363, p=0.007) while tortoises in the familiar condition did not (t_10_=1.077, p=0.303). These effects suggest in the stranger condition tortoises initially oriented more towards the experimental mate than familiar tortoises, while in the second part of the test tortoises in the stranger condition oriented less towards the other experimental subject than familiar tortoises.

**Figure 3.**
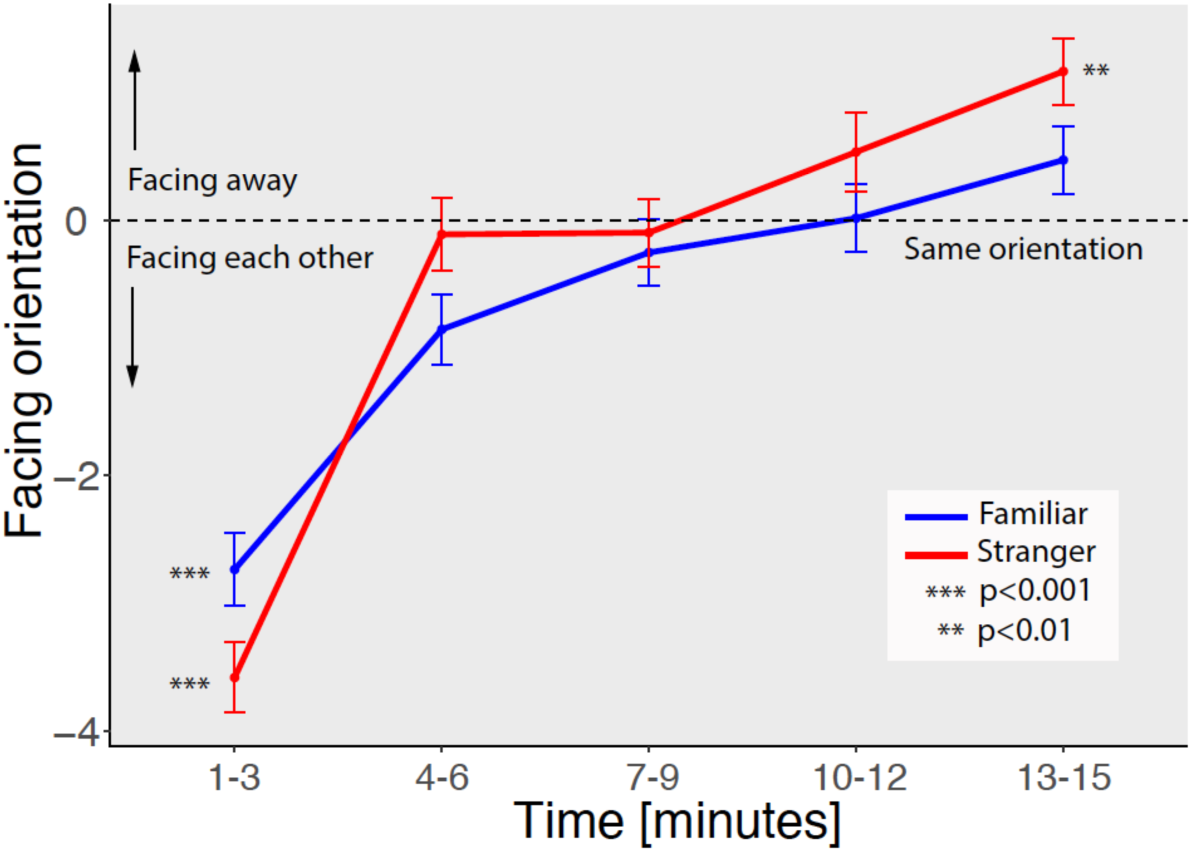
Facing orientation by Condition in Time. Negative values indicate tortoises facing each other, positive values opposite orientation. Significance against the chance level (same orientation, facing in the same direction) is indicated by asterisks: *** p<0.001, ** < 0.01.

## DISCUSSION

Individual recognition, the ability to recognize particular individuals, has been mainly investigated in contexts of repeated interactions, such as kin recognition, neighbour-stranger recognition, dominance hierarchies [reviewed in ,1,2]. Recognition of specific individuals is documented in different taxa, such as fish [16,17], mammals [18,19], reptiles [20,21], birds [22–24], invertebrates [25,26]. Since in most cases affiliative or competitive interactions recurred several times before recognition was tested, it is not clear to what extent individual recognition requires an individual to have experience with multiple subjects. The case of filial imprinting in precocial avian species is a notable exception: through this learning mechanism, chicks of the domestic fowl promptly recognize familiar individuals after a single exposure [3–6] and exhibit affiliative responses towards the imprinting stimulus. Chicks and other social species exhibit several predispositions for social behavioural soon after hatching [27], and these behaviours have a genetic component [28]. It is not known, though, whether exposure to a single individual is sufficient to elicit individual recognition at the onset of life in other species, including non-social animals. Showing abilities of individual recognition in tortoise hatchlings would suggest a certain independence of individual recognition from complex social experience and the possibility that this trait has evolved in contexts other than repeated social interactions.

Here, we documented the first evidence of spontaneous individual recognition in hatchlings of two species of tortoises (*Testudo graeca* and *Testudo marginata*) previously exposed to a single conspecific. At test, tortoises were placed in a novel environment, so that no territoriality was present for the test arena. Tortoises of both species showed strikingly different behavioural responses in the two conditions. While pairs of familiar tortoises appeared to ignore each other in the chosen trajectories, and progressively reached the chance level distance, pairs of stranger tortoises initially approached each other much faster than familiar pairs, then walked away from each other and reached a distance significantly larger than expected by chance. Interestingly, in the first part of the test unfamiliar individuals approached each other faster than familiar individuals, suggesting an initial interest in the unfamiliar conspecific. This observation is supported by the fact that tortoises in the unfamiliar condition faced more the other individual than tortoises in the familiar condition.

We hypothesize that the spontaneous avoidance of unfamiliar tortoises in hatchlings might be an adaptation to help dispersal of the clutch. In the wild, it has been documented that *T. marginata* and *T. graeca* lay multiple clutches per year (1-4) and each clutch contains 3-7 eggs [29,30] (in captivity, we observed larger clutch size, up to 12 eggs per individual in *T. marginata*). Hence, without active dispersal hatchlings would quickly saturate the carrying capacity of the environment, moreover, without dispersal there would be a greater exposure to predation of the entire clutch. In line with this idea, the range of hatchlings has little overlap [31] and females of *T. graeca* increase mobility before or after nesting, producing a wide dispersal of nests laid by each individual [32]. Overall, our results show that non-social species, such as land tortoises, possess the capacity for individual recognition at the onset of life, with very limited experience with other individuals. This suggests that individual recognition abilities might have evolved in contexts different than repeated social interactions.

## Authors’ contributions

E.V. and G.S conceived the project; G.S., E.V. and S.D. designed the experiment; S.D. carried out the experiment; E.V. M.C. and S.D. analysed the data; M.C. developed the simulations; E.V. drafted the paper; all authors revises the manuscript and gave final approval for publication.

## Acknowledgements

We thank the internship students of the high-school “Liceo A. Rosmini” (Rovereto, Italy) for help in data collection and the Rovereto Civic Museum Foundation, that provided the facilities to carry out the present research.

